# Combined single cell proteomics and transcriptomics reveals discrete human tendon cells populations persist in vitro and on fibrous scaffolds

**DOI:** 10.1101/2021.08.09.455617

**Authors:** Adrian Kendal, Antonina Lach, Pierre-Alexis Mouthuy, Rick Brown, Constantinos Loizou, Mark Rogers, Robert Sharp, Andrew Carr

## Abstract

Chronic tendinopathy represents a growing burden to healthcare services in an active and ageing global population. The ability to identify, isolate and interrogate, in vitro, key pathogenic and reparative tendon cell populations is essential to developing precision therapies and implantable materials.

Human hamstring tendon cells were cultured for 8 days on either tissue culture plastic or aligned electrospun fibres made of polydioxanone (absorbable polymer). Combined single cell surface proteomics and unbiased single cell transcriptomics (CITE-Seq) revealed six discrete cell clusters, four of which shared key gene expression determinants with ex vivo human cell clusters. These were PTX3_PAPPA, POST_SCX, DCN_LUM and ITGA7_NES cell clusters. Surface proteomics found that PTX3_PAPPA cells were CD10+CD26+CD54+. ITGA7_NES cells were CD146+, and POSTN_SCX cells were CD90+CD95+CD10+.

Three clusters preferentially survived and proliferated on the aligned electrospun fibres; DCN_LUM, POSTN_SCX, and PTX3_PAPPA. They maintained high expression of tendon matrix associated genes, including *COL1A1, COL1A2, COL3A1, ELN, FBLN1*, and up-regulated genesets enriched for TNF-*α* signalling via NFκB, IFN-Ɣ signalling and IL-6/ STAT3 signalling. When cells were pre-selected based on surface protein markers, a similar up-regulation of pro-inflammatory signalling pathways was observed, particularly in *PTX3* gene expressing CD10+CD26+CD54+ cells, with increased expression of genes associated with TNF-*α* signalling and IFN-γ signalling.

Discrete human tendon cell sub populations persist in vitro culture and can be recognised by specific gene and surface protein signatures. Aligned PDO fibres promote the survival of three clusters, including pro-inflammatory *PTX3* expressing CD10+CD26+CD54+ cells found in chronic tendon disease.

## Introduction

Musculoskeletal disorders are a major cause of long term morbidity, responsible for the second most years lived with disability worldwide (DALYs GBD 2015). Chronic tendinopathy affects over 20% of the population and the lower limbs are most commonly affected with an incidence of 18/1000 person years (de Jonge et al. 2011; Albers et al. 2016; Riel et al. 2019). At present there is no curative treatment for chronic tendon disease and patients are left to manage their symptoms with a combination of analgesics, lifestyle changes, physical therapy and ultimately surgical procedures with limited efficacy.

Our understanding of the cells and signalling pathways in diseased versus normal tendon is changing rapidly with the application of next generation sequencing techniques. It is only over the past three years that multiple discrete tendon cell sub populations have been described using single cell sequencing of both murine tendon (Tabula Muris et al. 2018; Giordani et al. 2019; Harvey, Flamenco, and Fan 2019), as well as healthy and diseased human tendon (Kendal AR 2020). Subsequent in situ transcriptomic analysis of human tendon tissue immediately ex vivo (Akbar et al. 2021) supported our original single cell sequencing atlas (Kendal AR 2020) and argued the case for pro-inflammatory stromal-immune cell interactions in shoulder tendinopathy. As burgeoning tendon cell atlases provide a greater insight into the pathogenesis of tendinopathy, it is likely that the ability to identify, isolate and interrogate tendon cells of interest in vitro will be fundamental in developing bespoke precision treatments.

There are a number of important challenges to overcome in achieving a reliable assessment of tenocyte behaviour in vitro. Firstly, tendon cells are sparsely distributed, particularly in healthy human tendon, and adherent to the dense surrounding extracellular matrix. This means that the cell yield can be low and necessitates mechanical and enzymatic processing that could influence cell characteristics. Secondly, the previously described tendon cell sub populations in mouse and human tendon, including most recently healthy versus diseased human tendon cells in vitro (Still C 2021), were based on single cell differential gene expression analysis. While there is some conservation between the species and overlap across different human tendon sites and different studies (Giordani et al. 2019; Kendal AR 2020; Akbar et al. 2021), it remains possible that the described clusters are a product of confounding transcriptomic variation. Moreover, the first application of CITE-Seq to human tendon identified only a small number of cell specific surface proteins and there is currently no optimal set of surface markers by which the various subsets can be identified (Kendal AR 2020). It is therefore unclear how well differentiated or pluripotent cells are within a given cluster or if cells across several clusters share a common progenitor. Finally, tendon cells have been shown to be mechano-sensitive, changing their shape, migration, proliferative rates and gene expression profile in response to differing culture surfaces (Lee et al. 2005; Bashur et al. 2009; Hakimi et al. 2015; Kendal et al. 2017; Still C 2021). It is likely that recreating the normal tendon architecture is a requisite condition for examining normal and diseased tendon cell behaviour in vitro. This is particularly important when developing implantable scaffolds aimed at improving tendon healing (Hakimi et al. 2015; Nezhentsev A 2021)

In order to begin to overcome some of these challenges, human hamstring tendon cells underwent 8 days of culture on either tissue culture plastic or electrospun polydioxanone (PDO) fibres. PDO has an established safety record as a suture material and can be electrospun into fibres to mimic tendon architecture (Mouthuy et al. 2015; Martins et al. 2020). CITE-Seq (Cellular Indexing of Transcriptomes and Epitopes by Sequencing) combined single cell surface proteomics and transcriptomics investigated whether different tendon cell clusters persist in vitro culture and if these clusters can be identified by surface protein markers.

The study identified multiple discrete tendon cell subtypes in vitro culture and identified those that preferentially proliferate on fibrous scaffolds. Differential gene expression and gene enrichment analysis revealed that TNF-*α*, IFN-γ, IL-6 and reparative matrix gene pathways were up-regulated by human tendon cells cultured on PDO electrospun fibre scaffolds.

## Materials and Methods

### Tendon collection and cell processing

Tendon biopsies were collected from six patients (average age of 37 years; 5 males, 1 female) with informed donor consent, in accordance with the Declaration of Helsinki, under ethics from the Oxford Musculoskeletal Biobank (09/H0606/11) and in compliance with National and Institutional ethical requirements. Only waste tissue that would otherwise have been disposed of was collected. Hamstring tendon was obtained from patients undergoing reconstruction of knee anterior cruciate ligament. Tendon samples measuring 10 (l) x 10 (w) mm were excised distal to the myotendinous junction and proximal to the enthesis. Samples were immediately placed in 4°C Iscove’s Modified Dulbecco’s Medium (IMDM) without antibiotics and without FCS. The tendons were rinsed in 1 x PBS, cut axially using a size 10 surgical scalpel into 1mm3 pieces and incubated at 37°C for 45 mins in Liberase (Merck) and 10ul/ml DNAse I (Thermo Scientific). Ham’s F-12 media + 10% FCS was added and the digested tissue passed through a 100μm cell strainer. Cells were then passaged to P1 with ‘culture media’ containing DMEM F12 media (Lonza, Slough, UK), 10% foetal bovine serum (Labtech, Uckfield, UK), and 1 % Penicillin-Streptomycin (Thermo Fisher Scientific, Waltham, MA, USA) at 37°C and 6% CO2. The culture medium was was replaced every 3 days.

### Fibrous scaffold preparation, cell seeding and cell culture

Polydioxanone (PDO) fibres were electrospun using a modified version of a previously described protocol (Hakimi et al. 2015). In brief, a 7 % w/v polymer solution of PDO (Riverpoint Medical, Portland, Oregon , USA) in 1,1,1,3,3,3-Hexafluoro- 2-propanol (HFIP, Halocarbon Product Corporation, North Augusta, South Carolina, USA) was prepared, with a compound that changes the conductivity of the solvent. Polymer solution was supplied by a syringe pump (integrated with IME electrospinning machine) at a flow rate of 0.8 mL/h. An IME electrospinner (IME Technologies, Spaarpot, The Netherlands) was used. PDO fibres were electrospun for up to 4 hours at 21°C, 30% relative humidity, from a double nozzle setup at voltage of 9.0 – 9.6 kV with a distance of 20.0 cm between the nozzle and the grounded collector. Aligned fibres of 800-1000nm in diameter were produced by electrospinning onto the collector – a drum covered with aluminium foil which was rotated at 2000RPM. Light microscopy confirmed a density 35 to 45 fibres per 0.01mm2. The electrospun PDO fibres were stored under vacuum desiccation for up to three months.

Aligned electrospun PDO fibres were stretched and secured over the surface of a 24-well plate Cell Crown™ insert (Sigma-Aldrich). The whole construct was sterilised in 100% ethanol for 60 mins and rinsed five times in sterile 1 x PBS. 5×10^5 (50μl of 1×10^6 cells/ml) human tendon cells suspended in culture media were seeded onto the PDO fibre CellCrown™ construct. Cells were allowed to adhere for 5 minutes before the CellCrown™ construct was inverted and firmly placed into a well of a 24 well plate containing culture media. This ensured that the all cells in the 24 well plate were attached to the PDO aligned fibres and not in contact with the tissue culture plastic surface. Every three days the culture medium was refreshed and each aligned PDO fibre CellCrown™ construct was transferred to a fresh 24 well plate. Technical triplets were performed for each biological sample. The control group consisted of 1×10^3 tendon cells, from the same donor, seeded directly to 24 well plates.

The tendon cells in both groups (the control tissue culture plastic group and the aligned PDO electrospun fibres group) were cultured in culture media containing DMEM F12 media + 10% foetal bovine serum + 1 % Penicillin/Streptomycin at 37°C and 6% CO2. The culture medium was replaced every 3 days and cells were cultured for a total of 8 days.

### Cell proliferation

Cell proliferation was quantified using AlamarBlue Assay, which has previously been validated (Ahmed et al., 1994; Voytik-Harbin et al., 1998). On day 1, day 4 and day 7 of culture, the PDO fibre CellCrown™ constructs were transferred into fresh 24 well plates to exclude from the analysis any cells no longer attached to the PDO fibres. 10% AlamarBlue (Invitrogen) in culture media was added to the fresh wells containing the PDO fibre CellCrown™ constructs as well as the 24 well plate containing cells cultured on tissue culture plastic. In both groups, the cells were incubated at 37°C for 4 hours. 100*μ*L samples (n = 3) were taken from each of the wells and pipetted into a 96-well plate. Fluorescence was measured using a FLUOstar Omega Microplate Reader (BMG Labtech, Aylesbury, UK) at 544 nm excitation and 590 nm emission and compared to a cell density calibration curve of hamstring tenocytes cultured on a 24 well plate under the same conditions. Technical triplicates were performed for each biological sample.

### CITE-seq

Cells were dissociated by incubation at 37°C for 10 minutes with 500μl per well of StemPro® Accutase® (ThermoFisher). Tendon cells cultured on tissue culture plastic or PDO electrospun fibres were washed and re-suspended in 100*μ*l staining buffer (1 x PBS + 2% BSA + 0.01% Tween). As per the CITE-Seq protocol (https://citeseq.files.wordpress.com/2019/02/cite-seq_and_hashing_protocol_190213.pdf), cells were then incubated for 10 minutes at 4°C in human Fc Blocking reagent (FcX, BioLegend). Cells were incubated at 4°C for a further 30 minutes with 0.5*μ*g of TotalSeq-A (Biolegend) monoclonal anti-CD10, anti-CD105, anti-CD146, anti-CD26, anti-CD31, anti-CD34, anti-CD44, anti-CD45, anti-CD54, anti-CD55, anti-CD90 (THY1), anti-CD95, anti-CD73, anti-CD9 and anti-CD140a antibodies. In addition, cells were incubated with 0.5*μ*g of one of eight surface hashing antibodies (Biolegend) so that their sample of origin could be identified following sequencing across two lanes as previously described (Kendal AR 2020).

The cells were then washed three times with staining buffer and re-suspended in 1 x PBS at 1000 cells/*μ*l. The cell suspensions were filtered using a 100*μ*m sieve. The final concentration, single cellularity and viability of the samples were confirmed using a haemocytometer. Cells were loaded into the Chromium controller (10x-Genomics) chip following the standard protocol for the Chromium single cell 3’ kit. A combined hashed cell concentration was used to obtain an expected number of captured cells between 5000-10000 cells. All subsequent steps were performed based on the CITE-Seq protocol (https://citeseq.files.wordpress.com/2019/02/cite-seq_and_hashing_protocol_190213.pdf). Libraries were pooled and sequenced across multiple Illumina HiSeq 4000 lanes to obtain a read depth of approximately 30,000 reads per cell for gene expression libraries.

The raw single cell sequencing data was mapped and quantified with the 10x Genomics Inc. software package CellRanger (v2.1) and the GRCh38 reference genome. Using the table of unique molecular identifiers produced by Cell Ranger, we identified droplets that contained cells using the call of functional droplets generated by Cell Ranger. After cell containing droplets were identified, gene expression matrices were first filtered to remove cells with > 5% mitochondrial genes, < 200 or > 5000 genes, and > 25000 UMI. Downstream analysis of Cellranger matrices was carried out using R (3.6.0) and the Seurat package (v 3.0.2, satijalab.org/seurat).

In total, 10,990 cells were selected for ongoing analysis after quality control filtering. Data were normalised using SCTransform function for RNA gene expression, hashed antibody (HTO) and surface antibody (ADT) expression level. Cells were selected based on high ‘expression level’ of their donor specific surface hashing antibody and low ‘expression level’ of the remaining hashing antibodies. Normalised data from all tendon cells were combined into one object and Integrated. Variable genes were discovered using the SCtransform function with default parameters. The FindIntegrationAnchors function command used default parameters (dims = 1:30) to discover integration anchors across all samples. The IntegrateData function was run on the Anchorset with default additional arguments. ScaleData and RunPCA were then performed on the integrated assay to compute 16 principal components (PC). Uniform Manifold Approximation and Projection (UMAP) dimensionality reduction was carried out and Shared Nearest Neighbour (SNN) graph constructed using dimensions 1:16 as input features and default PCA reduction (Becht et al. 2018). Clustering was performed on the Integrated assay at a resolution of 0.5 with otherwise default parameters( Butler et al. 2018).

Seurat FindAllMarkers was used to identify positive and negative markers of a single cluster compared to all other cells. Following expression level normalisation using SCTransform function, the average expression level of a feature (e.g. gene or surface protein) was calculated across each cluster. The minimum percentage of cells in which the feature is detected in the each group was set to 25% . The average log2 fold change threshold was set to at least 1.0 such that the average expression level of a feature would have to be at least double in one cluster compared to another to reach threshold. Significance was determined using Wilcoxon rank sum test with p-values adjusted based on bonferroni correction applying all features in the data set (p_val_adj <0.05).

Gene-set enrichment analyses were performed using R Bioconductor v3.13 package, with reference to Molecular Signatures Database v7.4 and Reactome Pathway Database.

## Results

### scRNA-seq reveals multiple tendon cell clusters in vitro

Healthy human hamstring tendon cells from six patient donors were split into two groups and cultured on either tissue culture plastic or electrospun aligned PDO fibrous scaffolds. After 8 days of culture, a total of 10,990 human hamstring tendon cells underwent single cell transcriptomic analysis, post quality control. 2,765 cells were cultured on PDO fibres and 8,225 cells were cultured on tissue culture plastic. Unsupervised graph based clustering and UMAP (uniform manifold approximation and projection (Becht et al. 2018)) of the integrated dataset (cells cultured on either tissue culture plastic or PDO fibres) revealed seven transcriptomic clusters (Figure 1). All clusters contained cells expressing high levels of genes associated with tenocytes including *COL1A1, COL1A2, COL3A1, FBLN, FBN1* and *ELN* and low expression of genes associated with endothelial, immune or skeletal muscle cells (Figure 1D). The top 15 differential expressed genes for each cluster are summarised in Figure 1E. PTX3_PAPPA and DCN_LUM clusters were predominantly composed of cells cultured on PDO aligned fibres. The majority of DKK1_FGF5 cells, PTX3_PAPPA cells, POSTN_SCX cells and PRELP_FMOD cluster cells were cultured on plastic (Figure 1).

**Figure 1.**
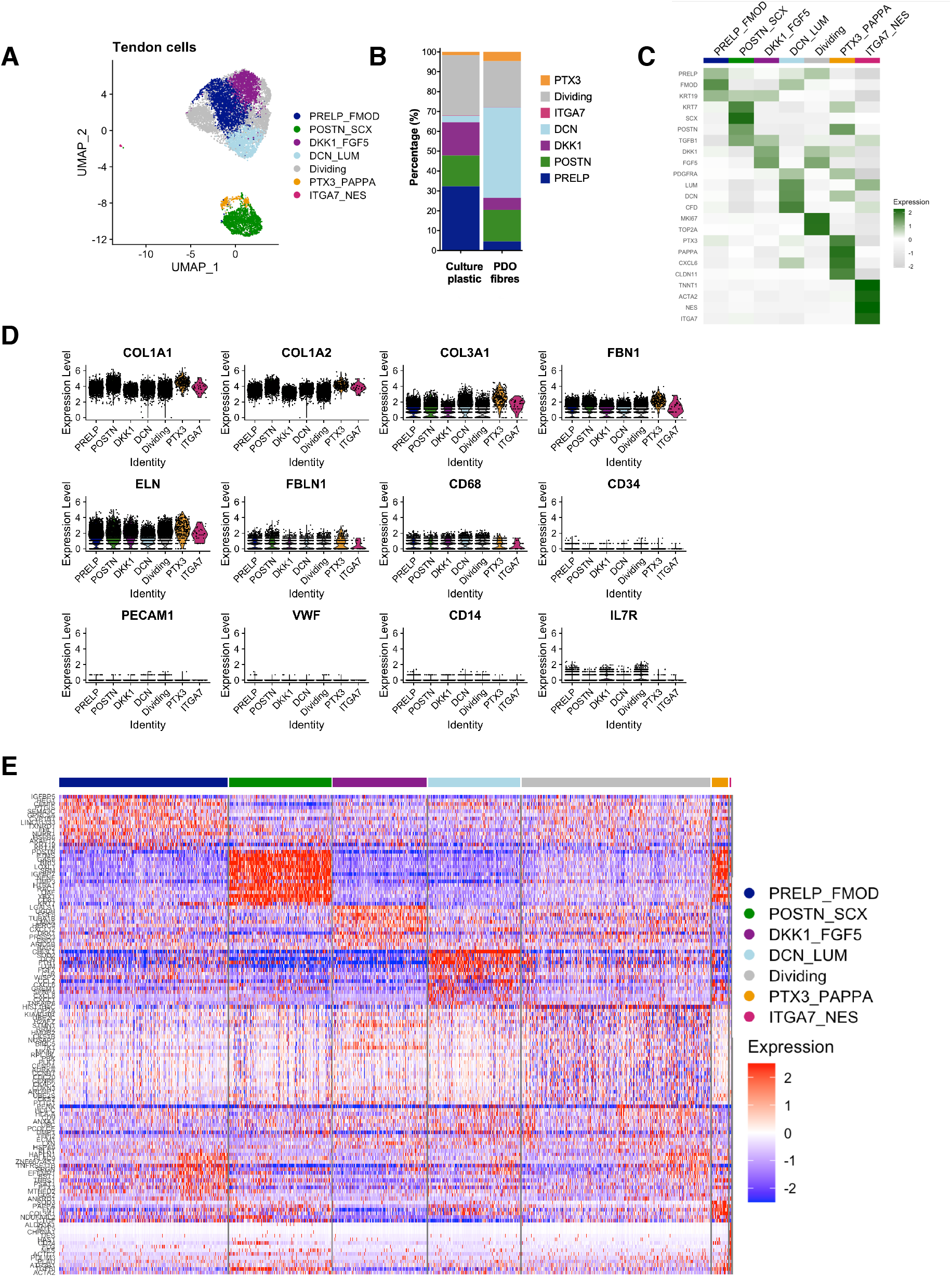
Single cell atlas of cultured human hamstring tendon cells. (A) Single cell transcriptomic UMAP dimensionality reduction of cultured human hamstring cells revealed six discrete cell clusters. The data represents 10,990 cells; 2,765 cultured on PDO fibres and 8,225 cells cultured on tissue culture plastic for 8 days. (B) Cluster percentages of cells cultured on tissue culture plastic and PDO fibres. (C) Heatmap of average expression of top three differentiating genes per cluster. (D) Violin plots show high expression of matrix genes *COL1A1, COL1A2, COL3A1, FBN1, ELN,* and *FBLN1* by all clusters. (E) Heatmap demonstrating relative expression level of top 15 genes for each cluster.

In order to help further define the clusters observed in vitro, expression of the top 20 differentiating genes previously found in the ex vivo human tendon clusters (Kendal AR 2020) was analysed for each in vitro cluster (Figure 2A). In vitro ITGA7_NES cells demonstrated increased expression of genes found in ex vivo ITGA7+ cells. In vitro PTX3_PAPPA cells demonstrated increased expression of genes found in ex vivo PTX3+ cells. In vitro DCN_LUM cells demonstrated increased expression of genes found in ex vivo APOD+ cells, and in vitro POSTN_SCX cells demonstrated increased expression of genes found in ex vivo KRT7/SCX+ cells (Figure 2A). Figure 2B demonstrates the top differentially expressed genes for each in vitro cluster with reference to the remaining clusters (labelled genes indicating those expressed at >0.5 Log2Fold change, p<0.001).

**Figure 2.**
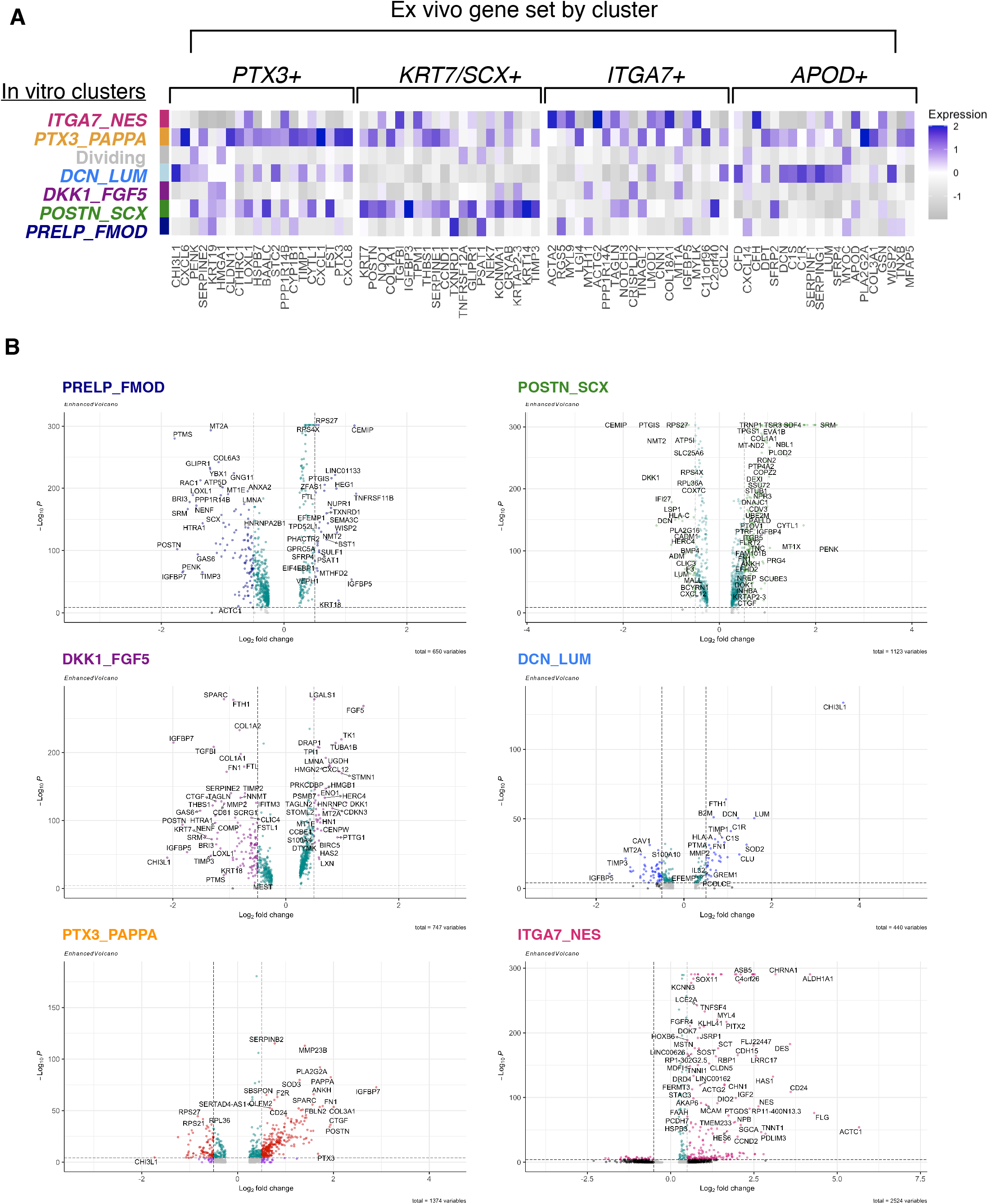
Differential gene expression of hamstring cells cultured in vitro. The data represents the integrated set of 10,990 cells cultured on either tissue culture plastic or aligned PDO fibres. (A) Heatmap showing the relative average expression for each in vitro cluster (y axis) of the top 20 genes previously found to be expressed by ex vivo human tendon cell clusters (x axis, historical dataset). (B) Volcano plots of differential genes expression for each in vitro cluster (labelled genes >0.5 Log2fold change in expression, p<0.001).

CITE-Seq surface proteomic analysis was performed using Total-SeqA (Biolegend) oligo-nucleotide conjugated monoclonal antibodies against twelve cell surface protein markers (Figure 3). This revealed that PTX3_PAPPA cells expressed high levels of surface CD10, CD26 and CD54 proteins. CD146 protein was up-regulated on the cell surface of ITGA7_NES cluster cells. POSTN_SCX cells were CD90+CD95+CD10+. PRELP_FMOD cells were CD90^low^CD10^neg^. Low surface expression of CD10, CD26 and CD54 protein was observed on cells in the DCN_LUM cluster. DKK1_FGF5 cluster cells did not express high levels of any of the surface proteins targeted in this study.

**Figure 3.**
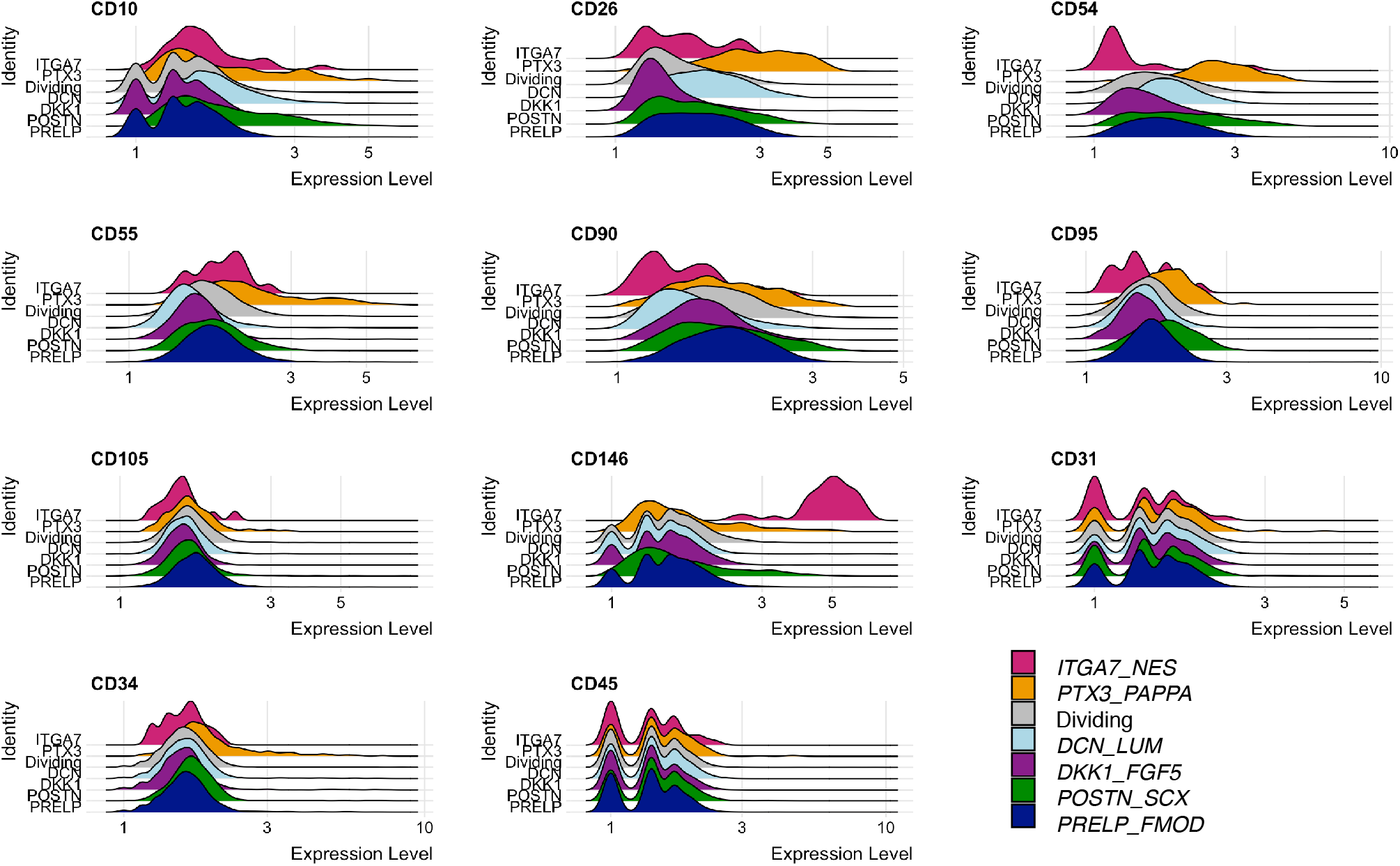
CITE-seq differential surface protein expression of hamstring cells cultured in vitro. The data represents the integrated set of 10,990 cells cultured on either tissue culture plastic or aligned PDO fibres. Ridgeplots of surface proteins detected by oligo-nucleotide conjugated anti-CD10, anti-CD26, anti-CD54, anti-CD55, anti-CD90, anti-CD95, anti-CD105, anti-CD31, anti-CD34, anti-CD146, and anti-CD45 for each cell cluster of the integrated dataset.

### Characteristics of cells preferentially cultured on aligned PDO electrospun fibres

The cells grown on PDO electrospun fibres proliferated but at a significantly lower rate over 8 days compared to cells cultured on plastic (Figure 4A, mean 1.7 fold increase versus 5.1 fold increase in cell numbers respectively, p<0.001). There was no significant difference in expression of *COL1A1, COL1A2, COL3A1, FBN1, ELN* and *FBLN1* between cells cultured on PDO fibres versus tissue culture plastic (Figure 4B, grey versus white respectively). Split violin plots demonstrate that comparable levels of surface CD10, CD26, CD54, CD90, CD95 and CD105 were seen for cells culture on plastic and electrospun PDO fibres across all clusters (Figure 4C). CD146 surface protein expression was greater on ITGA7_NES cluster cells cultured on plastic compared to PDO electrospun fibres, but very few ITGA7_NES cells survived on PDO fibres (Figures 1B, 4C).

**Figure 4.**
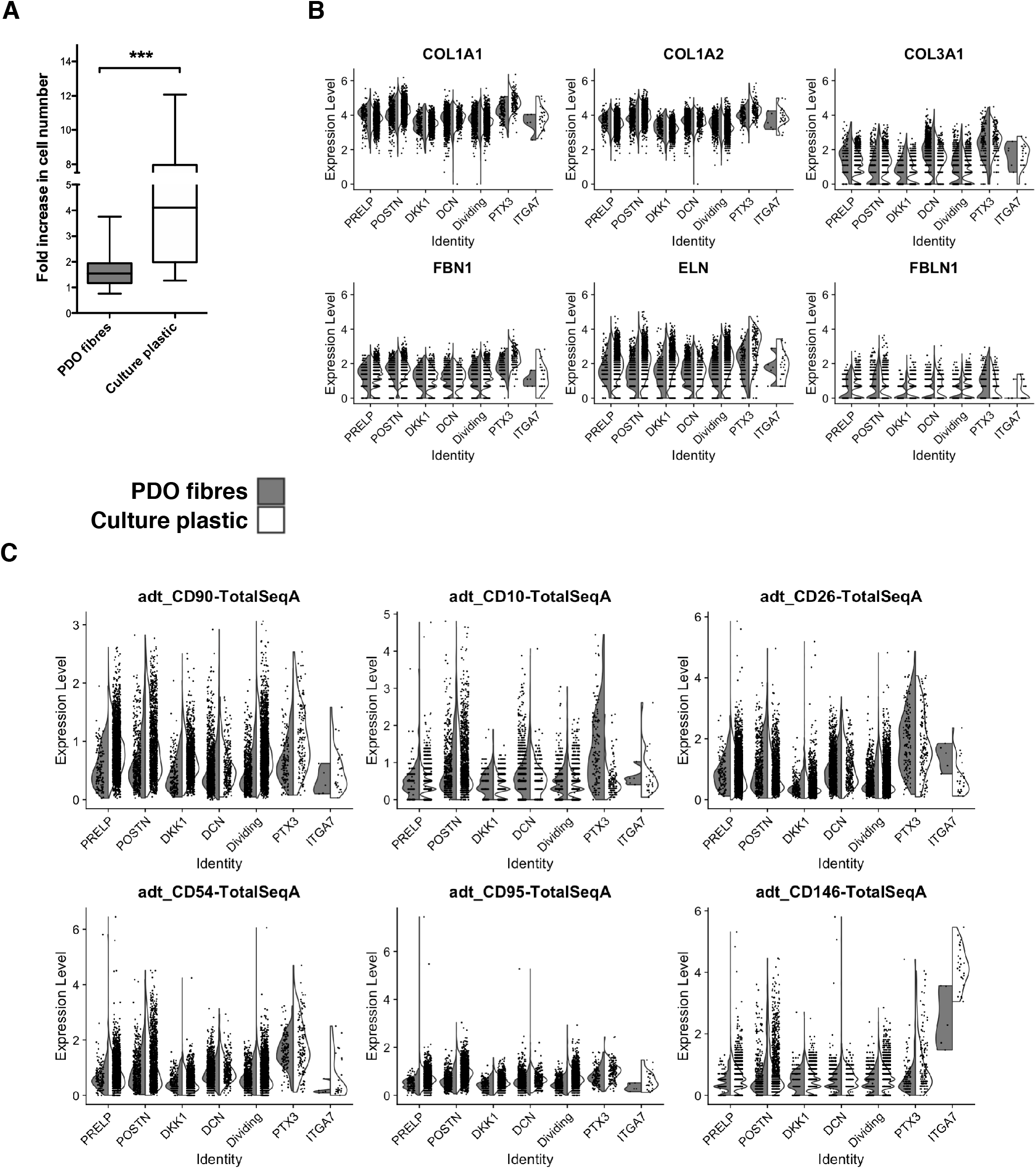
Human tendon cells cultured on tissue culture plastic vs PDO fibres. (A) Cells cultured on tissue culture plastic proliferated at significantly greater rate than those on PDO fibres (5.1 versus 1.7 fold increase over 7 days respectively, p<0.001). (B) Split violin plots showed there was no significant difference in expression of matrix genes *COL1A1, COL1A2, COL3A1, FBN1, ELN* and *FBLN1* between cells on PDO fibres (grey) and tissue culture plastic (white). (C) Split violin plots of oligo-nucleotide conjugated mAb recognising surface proteins show no significant difference in surface CD90, CD10, CD26, CD54, and CD95 between cells on PDO fibres (grey) versus tissue culture plastic (white). Very few ITGA7_NES cells survived on PDO fibres and had reduced surface CD147.

DCN_LUM, POSTN_SCX and PTX3_PAPPA cells preferentially survived on PDO electrospun scaffolds (Figure1B). In these three clusters, significantly greater expression of *SOD2, CXCL1*, *CXCL6*, and *CXCL8* was observed in cells cultured on PDO electrospun fibres versus tissue culture plastic (log2 fold change >0.5, p <0.05; Figure 5A). *COL6A3* was significantly increased in cells from PRELP_FMOD, DCN_LUM, PTX3_PAPPA, and DKK1_FGF5 when cultured on PDO electrospun fibres versus tissue culture plastic. *COL3A1* expression was increased in DCN_LUM cells and PRELP_FMOD cells cultured on PDO electrospun fibres. PRELP_FMOD cells additionally demonstrated increased expression of *COL8A1*, *DCN*, *FN1*, *LUM* and *MMP2* when cultured on PDO electrospun fibres.

**Figure 5.**
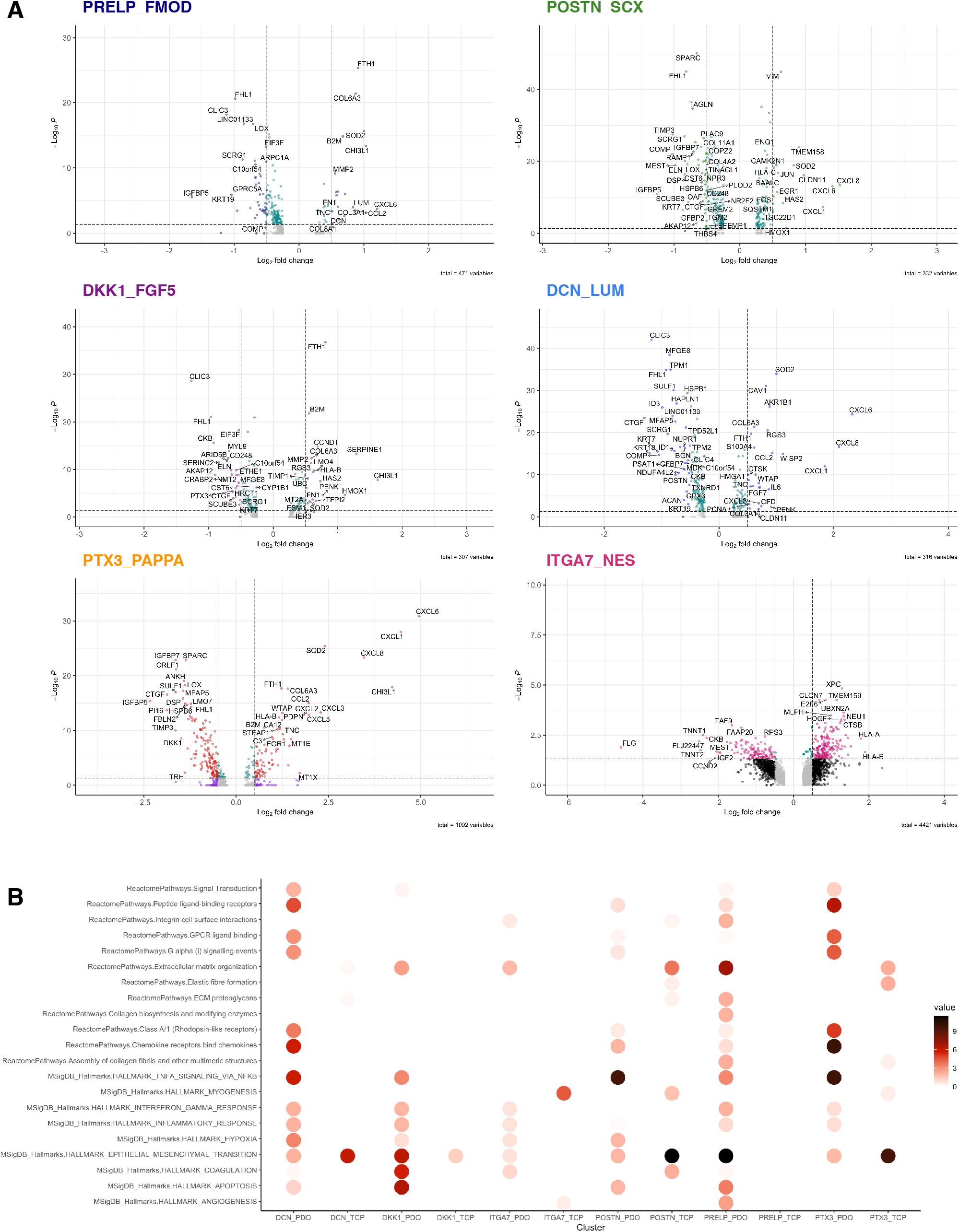
Differential gene expression of hamstring cells cultured on PDO fibres versus culture plastic. (A) Volcano plots of differential gene expression on PDO fibres versus tissue culture plastic for each in vitro cluster (labelled genes >0.5 Log2fold change in expression, p<0.001). (B) GeneOntology enrichment analysis of gene sets up-regulated by each cluster when cultured on PDO fibres (_PDO) versus tissue culture plastic (_TCP labelled x axis) with reference to Reactome Pathway and MSigDB HALLMARK databases (y axis).

Gene Ontology mapping was undertaken for differentially expressed genes observed in cells cultured on electrospun PDO fibres compared to tissue culture plastic (Figure 5B). Hallmark gene set enrichment analysis was performed with reference to the Molecular Signatures Database of 50 well defined biological processes and the Reactome Pathway Database. Culturing human hamstring tendon cells on PDO electrospun fibres up-regulated gene sets associated with ‘TNF-*α* signalling via NFκB’ and the ‘Inflammatory Response’ in all clusters. An increase in genes associated with ‘IFN-γ signalling’ was observed for cells cultured on PDO electrospun fibres from all clusters except POSTN_SCX. IL-6/STAT3 signalling was increased in POSTN_SCX, DCN_LUM and PTX3_PAPPA clusters (Figure 5B). A corresponding reduction in expression of genes associated with ‘Hallmark Myogenesis’ was observed in ITGA7_NES, POSTN_SCX and PTX3_PAPPA cluster cells cultured on PDO electrospun fibres compared with those from tissue culture plastic alone.

### Cell selection based on surface protein expression

CITE-Seq single cell surface proteomics allowed cells to be selected from the integrated dataset based on surface protein expression *before* performing further single cell gene expression analysis. In this way, cells were segregated into groups based on high levels of expression of surface protein CD10, CD26, CD54, CD90, CD95, CD105 or CD146. Differential gene expression of cells cultured on PDO electrospun fibres versus plastic were analysed for each group (Figure 6). An increase in TNF-*α* signalling gene sets was observed for CD26+ cells, as well as CD54+, CD90+, CD95+, CD105+ and CD146+ cells cultured on PDO electrospun fibres (Figure 6). CD10+ cells cultured on PDO electrospun fibres up-regulated gene sets associated with ‘INF-Ɣ response’, ‘TNF-*α* signalling’, and ‘Chemokine receptor binding chemokines’ (Figure 6E). In comparison, gene sets associated with myogenesis were down-regulated versus cells cultured on plastic alone. ‘Hallmark IL-6_JAK_STAT’ signalling gene expression was increased in CD54+ cells and CD90+ cells on electrospun PDO fibres. CD90+ cells, CD95+ cells and CD105+ cells cultured on PDO fibres up-regulated genes associated with IFN-Ɣ signalling. Gene sets associated with extra cellular matrix production pathways, including ‘Epithelial to mesenchymal transition’, ‘Extra cellular matrix organisation’, ‘Collagen formation’, were increased in CD26+ cells, CD54+ cells, CD90+ cells, CD95+ cells and CD105+ cells when cultured on PDO fibres. In contrast, CD146+ cells cultured on electrospun PDO fibres down-regulated ‘Epithelial to mesenchymal transition’ and ‘Extra cellular matrix organisation’ gene sets compared to those cultured on tissue culture plastic (Figure 6B).

**Figure 6.**
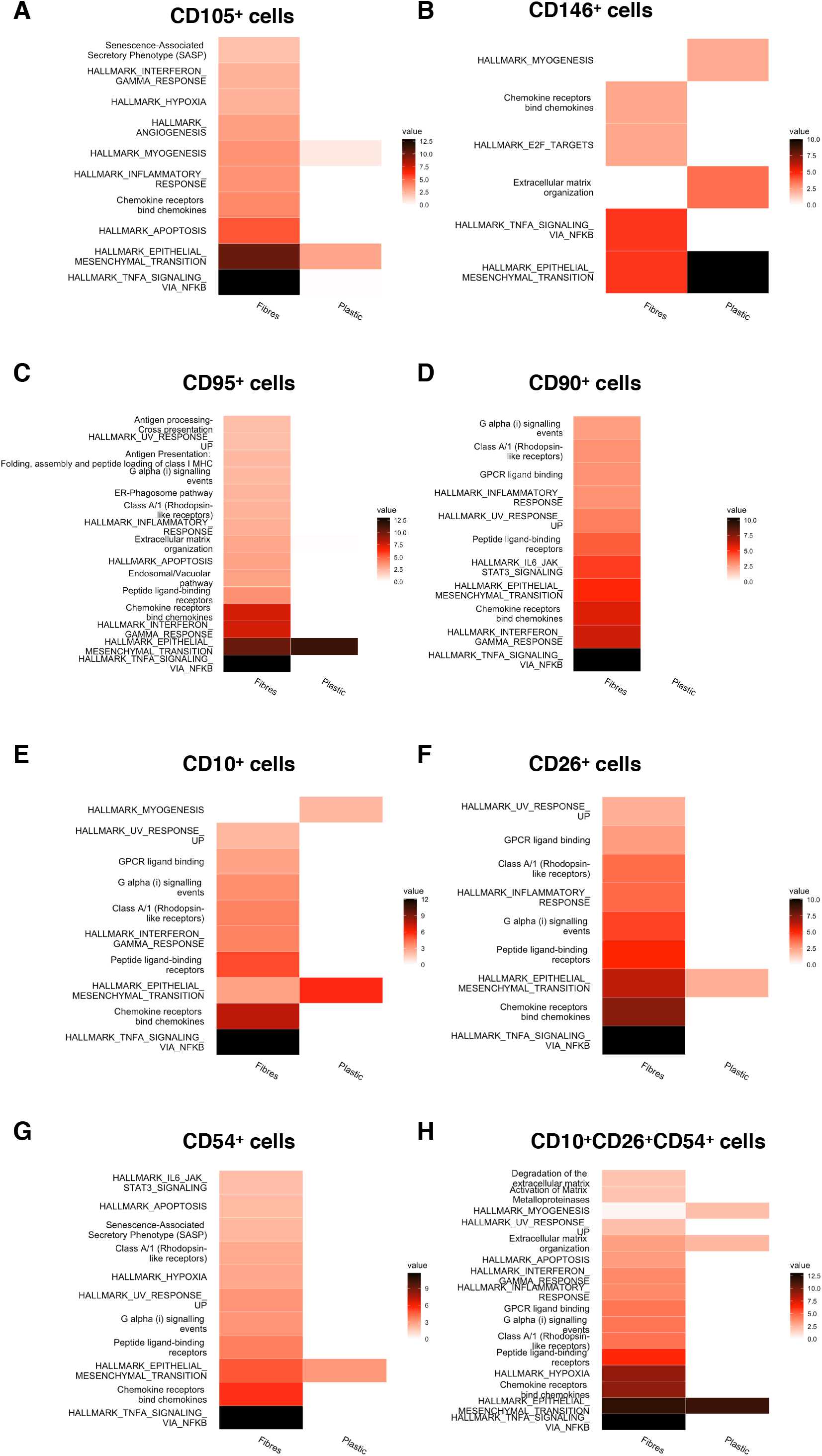
GeneOntology enrichment analysis of pre-selected hamstring cells cultured on PDO fibres versus culture plastic. Tendon cells were selected based on surface expression of (A) CD105, (B) CD146, (C) CD95, (D) CD90, (E) CD10, (F) CD26, (G) CD54 and (H) CD10 + CD26 + CD54 + proteins from the combined in vitro dataset. Gene Ontology enrichment analysis of differential gene expression was performed for each selected group of cells cultured on PDO fibres versus tissue culture plastic. Reactome Pathway and MSigDB HALLMARK databases were used for reference.

Cells that were selected based on combined surface expression of CD10+CD26+CD54+ exhibited increased expression of genes enriched for ‘Hallmark TNF-*α* signalling’, ‘INF-Ɣ response’, and ‘Inflammatory response’ when cultured on aligned PDO fibres. They also up-regulated genes associated with ‘Chemokine receptors bind chemokines’, ‘Degradation of extracellular matrix’, ‘Activation of Matrix Metalloproteinases’ and ‘Extracellular matrix organisation’ (Figure 6H). Genes enriched for ‘Hallmark myogenesis’ were down-regulated on culture with aligned PDO fibres versus tissue culture plastic.

## Discussion

Single cell RNA sequencing of human hamstring tendon cells has shown that multiple discrete cell clusters persist in vitro culture on both tissue culture plastic and aligned PDO electrospun fibres. There were six main sub populations including PRELP_FMOD, POSTN_KRT7, DKK1_FGF5, DCN_LUM, PTX_PAPPA and ITGA7_NES cells. All cell clusters expressed high levels of *COL1A1, COL1A2, COL3A1, ELN, FBLN1* genes associated with normal tendon matrix proteome (Hakimi et al. 2017). In comparison to initial (Kendal AR 2019, 2020) and subsequent (Akbar et al. 2021) ex vivo human tendon single cell atlases, endothelial cells, monocytes and lymphocytes were not present after 8 days of culture under conditions that favour tendon fibroblasts.

Four of the in vitro cell clusters shared gene expression profiles with ex vivo cell populations from a previous human tendon atlas (Kendal AR 2020) and Figure 2A). These were in vitro POSTN_KRT7, PTX3_PAPPA, ITGA7_NES and DCN_LUM clusters that showed increased expression of the top differentially expressed genes originally found in the ex vivo POSTN+, PTX3+, ITGA7+ and APOD+ clusters respectively. This supports the possibility that human tendon cell clusters represent populations that can be isolated ex vivo and remain discrete subtypes in vitro (Still C 2021). There was little in the gene expression profile of cultured DKK1_FGF5 and PRELP_FMOD to relate them to cell clusters found in the ex vivo atlas. These two clusters may represent confounding variation, particularly batch variation, or represents transient states of less well differentiated tendon cells that are selected by particular in vitro culture conditions.

The ability to identify cell populations by specific surface markers is essential to future isolation of live cells from each cluster (Cappellesso-Fleury et al., 2010; Halfon et al., 2011; Mohanty et al., 2014). CITE_seq proteomics revealed that surface markers for two in vitro clusters, ITGA7_NES cells and PTX3_PAPPA, were consistent with previous ex vivo findings. Human ITGA7+ cells ex vivo had high levels of CD146 protein surface expression (Kendal AR 2020), as did the cluster of in vitro ITGA7_NES cells (Figure 3). PTX3+ cells were CD10+CD26+CD54+ ex vivo (Kendal AR 2020), and PTX3_PAPPA cells in this dataset were also CD10+CD26+CD54+ whether cultured on tissue culture plastic or PDO electrospun fibres. This was not the case for the other two clusters; POSTN_KRT7 cells in vitro were CD90+CD95+CD10+ whereas POSTN+ ex vivo cells were CD90+CD105+CD146+ and DCN_LUM cluster cells in vitro had low surface expression of CD10, CD26 and CD54 protein compared to APOD+ cells ex vivo that were CD90+CD34+ (Figure 3).

ITGA7_NES cells were first described in murine muscle as smooth muscle-mesenchymal cells situated in perivascular regions (Yin et al. 2016; Giordani et al. 2019). They seem to reproducibly express *ACTA2*, *RGS5*, *MYL9*, and *TAGLN* in murine tendon cells ex vivo (Yin et al. 2016; Giordani et al. 2019) as well as in human tendon cells ex vivo (Kendal AR 2020) and, now in vitro. As with previous studies in which ITGA7+ (Kendal AR 2020) or *NES*+ cells expanded on tissue culture plastic (Yin et al. 2016), we found ITGA7_NES cells cells preferentially grew on tissue culture plastic and very few survived on PDO electrospun fibres (Figure 1B). Those that did had reduced expression of Hallmark genes associated with myogenesis and an increase in ‘Epithelial to Mesenchymal Transition’ genes and ‘Interferon Gamma Response’ genes (Figure 5).

PTX3+ cells from chronic diseased tendon have previously been shown ex vivo to up-regulate *CHI3L*, *CLDN11*, *PENK*, *SERPINE2* and pro-inflammatory genes *CXCL1*, *CXCL6* and *CXCL8,* suggesting they may a play a role in driving inflammation in chronic tendinopathy (Kendal AR 2020). PTX3_PAPPA cells preferentially survived and proliferated on aligned PDO electrospun fibres (Figure 1B) and again increased expression of *CHI3L, CXCL1, CXCL3, CXCL6,* and *CXCL8* (Figure 5). Similarly, selecting *PTX3* expressing cultured cells based on combined surface expression of CD10+CD26+CD54+ showed increased expression of genes associated with TNF-*α* signalling, INF-γ signalling and G-coupled chemokine signalling pathways (Figure 6H). The increased expression of “Matrix Metalloproteinases’ genes observed in CD10+CD26+CD54+ cells on aligned PDO fibres suggests that *PTX3*+ cells may be involved in the early response to tendon disruption; recruiting immune cells and degrading abnormal tendon matrix to perform the groundwork for the next phases of tendon repair.

It is unclear from these very early findings if *PTX3* expressing tenocytes are responding to a loss of tendon homeostasis by up-regulating pro-inflammatory genes in an attempt to resolve micro-structural damage or if they are inappropriately driving chronic inflammation. In either case these results suggest that aligned electrospun PDO fibres provide a useful scaffold and environment for their ongoing investigation in vitro. A similar population of cells expressing *IL8*, *CXCL1/6/8* and *PTX3* were described in single cell transcriptomics of healthy and diseased patellar tendon progenitor cells cultured under mechanical stress (Still C 2021). The majority of these named ‘pro-inflammatory Tendon Progenitor Cells’ were from diseased human tendon and expressed gene sets enriched for pro-inflammatory signalling pathways including ‘Interleukin-1 regulation of extracellular matrix’. These complimentary ex vivo and now in vitro findings add to the increasing evidence in favour of pro-inflammatory tendon fibroblasts driving chronic tendon disease, possibly in response to a loss of normal tendon homeostasis (Dakin et al. 2018; Akbar et al. 2021). The ability to focus on a particular tendon cell subsets will greatly advance our understanding of how tenocytes interact with immune cells (Stolk et al. 2017; Garcia-Melchor et al. 2021). We now have the means to isolate (based on CD10+CD26+CD54+ surface markers) and the means to culture (PDO aligned fibres) *PTX3+* cells implicated in chronic human tendinopathy.

In general cells proliferated at a slower rate on aligned PDO fibres compared to tissue culture plastic. This is consistent with previous observations (Kendal et al. 2017). Tendon fibroblast undergo a dramatic change in their morphology when seeded onto electrospun fibres, spreading along and across aligned fibres and using them as a scaffold on which to migrate( Hakimi et al. 2015; Kendal et al. 2017). The mechano-sensitivity of tendon cells to their surface environment is not restricted to electrospun fibres and is well documented on multiple structures (Lee et al. 2005; Bashur et al. 2009; Kim et al. 2009; Fleischer et al. 2015; Gomes et al. 2015; Smith et al. 2016). Historically it has not been clear which tendon cells are more likely to adhere to the fibres, which preferentially survive and whether there are transcriptional and phenotypic differences in the responses of different subtypes. In this study, three cell clusters preferentially survived on PDO electrospun fibres; DCN_LUM, PTX3_PAPPA and POSTN_SCX clusters (Figure 1B). These cells proliferated on PDO electrospun fibres and continued to express *COL1A1, COL1A2, COL3A1, FBN1, FBLN1* and *ELN* at similar levels to cells cultured on tissue culture plastic (Figure 4B). Gene enrichment analysis revealed an up-regulation of Hallmark gene sets for ‘TNF-*α* signalling via NFκB’ and ‘IL6_JAK_STAT6′ signalling in all three clusters (Figure 5B). DCN_LUM and PTX3_PAPPA cells up-regulated Hallmark genes sets for ‘IFN-Ɣ signalling’. In all three clusters, there was a relative reduction in expression of genes associated with epithelial to mesenchymal transition.

A similar up-regulation in IL6-JAK-STAT3 signalling and TNF-α signalling via NF-κB was observed when hamstring tendon cells were cultured for 14 days on twisted electrospun PDO compared to tissue culture plastic and smooth PDS II (polydioxanone suture material) (Nezhentsev A 2021). Gene enrichment for Mtorc 1 signalling was also up-regulated, while gene sets associated with epithelial-to-mesenchymal transition were down-regulated. Again, it is not clear if the observed pro-inflammatory signalling is a desirable early response to tendon damage and represents a favourable state when designing and screening implantable scaffolds to treat chronic tendinopathy.

In the absence of defined surface markers by which cells of interest can be manually selected in vitro, CITE-Seq allows transcriptomic analysis of cells ‘virtually’ pre-selected based on high expression of a given surface marker. Split Violin graphs of surface protein expression demonstrate that there was no significant decrease in surface CD90, CD10, CD26, CD54, CD95 of cells cultured on aligned PDO electrospun fibres versus tissue culture plastic (Figure 4C). We could therefore select cells from the integrated dataset that have high surface expression of these proteins. Genes associated with inflammatory response and/or pro-inflammatory signalling pathways were seen for CD90, CD10, CD26, CD54, CD95 and CD105 cells cultured on PDO fibres (Figure 6). As before, CD146+ cells (which are predominantly ITGA7_NES cells) down-regulated Hallmark genes enriched for ‘Myogenesis’, ‘Extra-cellular matrix organisation’ and ‘Epithelial to mesenchymal transition’ (Figure 6G).

We focused on a small set of surface proteins and future studies will greatly benefit from expanding the repertoire. We are still not in a position to identify and isolate tendon cell populations of interest with the notable exceptions of immune cells, endothelial cells, *ITGA7/NES* and *PTX3* expressing cells. This study was limited to cells from one type of tendon, a short period of in vitro culture and only two different culture conditions. It is still not clear how to define normal tendon cell behaviour in vitro and which culture conditions are optimal. Finally, batch variation is a major limitation in single cell RNA sequencing of cells cultured in vitro and emphasises the importance of validating any transcriptomic descriptive findings.

Our results demonstrate that discrete cell clusters can be identified by specific gene and surface protein signatures when human hamstring tendon cells are cultured in vitro. The presence of multiple different types of human tendon cell, and their persistence in culture, question the relevance of ongoing investigations that rely on the pooled responses of unsorted tendon cells. Further work is required to identify and sort tendon cell subsets, for example PTX3_PAPPA cells were found to be CD10+CD26+CD54+, but no clear surface markers were found for DCN_LUM cells based on the very limited set of monoclonal antibodies used in this study. It remains to be seen whether there is any functional relevance to the described cell clusters, and ongoing in vitro interrogation of subset behaviour and phenotypic stability is likely to require consideration of culture surfaces; PTX3_PAPPA cells proliferated on aligned PDO fibres while ITGA7_NES cells preferred tissue culture plastic.

## Conclusion

Combined single cell transcriptomics and proteomics of human hamstring tendon cells demonstrated multiple discrete clusters that persist in vitro culture. Four cell clusters closely resembled ex vivo human tendon cell clusters in their gene expression profile; PTX3_PAPPA, ITGA7_NES, DCN_LUM and POSTN_KRT7 cells. Culture on aligned PDO electrospun fibres favoured DCN_LUM, POSTN_SCX and PTX3_PAPPA cells, maintaining expression of common tendon matrix genes and up-regulating gene sets enriched for inflammatory signalling.

*PTX3* expressing cells have been implicated in chronic tendon disease. By demonstrating that they are CD10+CD26+CD54+, proliferate on aligned PDO fibres and up-regulate gene sets associated with TNF-*α* and IFN-γ, we have provided an opportunity to interrogate further these disease associated cells in vitro.

## Acknowledgements

Hubert Slawinski, Theo Kyriakou, Santiago Revale and Rory Bowden (Oxford Genomics Centre)

There are no competing interests.

## Notes

There are no conflicts of interest

### Competing Interest Statement

The authors have declared no competing interest.

## References

Akbar, M., MacDonald, L., Crowe, L. A. N., Carlberg, K., Kurowska-Stolarska, M., Stahl, P. L., Snelling, S. J. B., McInnes, I. B., and Millar, N. L. (2021), “Single cell and spatial transcriptomics in human tendon disease indicate dysregulated immune homeostasis,” Ann Rheum Dis. DOI: 10.1136/annrheumdis-2021-220256.

Albers, I. S., Zwerver, J., Diercks, R. L., Dekker, J. H., and Van den Akker-Scheek, I. (2016), “Incidence and prevalence of lower extremity tendinopathy in a Dutch general practice population: a cross sectional study,” Bmc Musculoskeletal Disorders, 17.

Bashur, C. A., Shaffer, R. D., Dahlgren, L. A., Guelcher, S. A., and Goldstein, A. S. (2009), “Effect of fiber diameter and alignment of electrospun polyurethane meshes on mesenchymal progenitor cells,” Tissue Eng Part A, 15 (9), 2435–2445. DOI: 10.1089/ten.tea.2008.0295.

Becht, E., McInnes, L., Healy, J., Dutertre, C. A., Kwok, I. W. H., Ng, L. G., Ginhoux, F., and Newell, E. W. (2018), “Dimensionality reduction for visualizing single-cell data using UMAP,” Nat Biotechnol.

Butler, A., Hoffman, P., Smibert, P., Papalexi, E., and Satija, R. (2018), “Integrating single-cell transcriptomic data across different conditions, technologies, and species,” Nat Biotechnol, 36 (5), 411–420. DOI: 10.1038/nbt.4096.

Dakin, S. G., Newton, J., Martinez, F. O., Hedley, R., Gwilym, S., Jones, N., Reid, H. A. B., Wood, S., Wells, G., Appleton, L., Wheway, K., Watkins, B., and Carr, A. J. (2018), “Chronic inflammation is a feature of Achilles tendinopathy and rupture,” Br J Sports Med, 52 (6), 359–367.

DALYs GBD, C. G. B. D. R. F. C. (2015), “Global, regional, and national comparative risk assessment of 79 behavioural, environmental and occupational, and metabolic risks or clusters of risks in 188 countries, 1990-2013: a systematic analysis for the Global Burden of Disease Study 2013,” Lancet, 386 (10010), 2287–2323.

de Jonge, S., van den Berg, C., de Vos, R. J., van der Heide, H. J. L., Weir, A., Verhaar, J. A. N., Bierma-Zeinstra, S. M. A., and Tol, J. L. (2011), “Incidence of midportion Achilles tendinopathy in the general population,” British Journal of Sports Medicine, 45 (13), 1026–1028. DOI: 10.1136/bjsports-2011-090342.

Fleischer, S., Miller, J., Hurowitz, H., Shapira, A., and Dvir, T. (2015), “Effect of fiber diameter on the assembly of functional 3D cardiac patches,” Nanotechnology, 26 (29), 291002. DOI: 10.1088/0957-4484/26/29/291002.

Garcia-Melchor, E., Cafaro, G., MacDonald, L., Crowe, L. A. N., Sood, S., McLean, M., Fazzi, U. G., McInnes, I. B., Akbar, M., and Millar, N. L. (2021), “Novel self-amplificatory loop between T cells and tenocytes as a driver of chronicity in tendon disease,” Ann Rheum Dis. DOI: 10.1136/annrheumdis-2020-219335.

Giordani, L., He, G. J., Negroni, E., Sakai, H., Law, J. Y. C., Siu, M. M., Wan, R., Corneau, A., Tajbakhsh, S., Cheung, T. H., and Le Grand, F. (2019), “High-Dimensional Single-Cell Cartography Reveals Novel Skeletal Muscle-Resident Cell Populations,” Mol Cell, 74 (3), 609–621 e606.

Gomes, S. R., Rodrigues, G., Martins, G. G., Roberto, M. A., Mafra, M., Henriques, C. M., and Silva, J. C. (2015), “In vitro and in vivo evaluation of electrospun nanofibers of PCL, chitosan and gelatin: a comparative study,” Mater Sci Eng C Mater Biol Appl, 46, 348–358. DOI: 10.1016/j.msec.2014.10.051.

Hakimi, O., Mouthuy, P. A., Zargar, N., Lostis, E., Morrey, M., and Carr, A. (2015), “A layered electrospun and woven surgical scaffold to enhance endogenous tendon repair,” Acta Biomaterialia, 26, 124–135. DOI: 10.1016/j.actbio.2015.08.007.

Hakimi, O., Ternette, N., Murphy, R., Kessler, B. M., and Carr, A. (2017), “A quantitative label-free analysis of the extracellular proteome of human supraspinatus tendon reveals damage to the pericellular and elastic fibre niches in torn and aged tissue,” PLoS One, 12 (5), e0177656.

Harvey, T., Flamenco, S., and Fan, C. M. (2019), “A Tppp3(+)Pdgfra(+) tendon stem cell population contributes to regeneration and reveals a shared role for PDGF signalling in regeneration and fibrosis,” Nat Cell Biol.

Kendal, A., Snelling, S., Dakin, S., Stace, E., Mouthuy, P. A., and Carr, A. (2017), “Resorbable electrospun polydioxanone fibres modify the behaviour of cells from both healthy and diseased human tendons,” Eur Cell Mater, 33, 169–182.

Kendal AR, L. T., Al-Mossawi H, Appleton L, Dakin S, Brown R, Loizou C, Rogers M, Sharp R, Carr A. (2020), “Multi-omic single cell analysis resolves novel stromal cell populations in healthy and diseased human tendon.,” Sci Rep., 3 (10). DOI: 10.1038/s41598-020-70786-5.

Kendal AR, L. T., Al-Mossawi H, Brown R, Loizou C, Rogers M, Sharp M, Dakin S, Appleton L, Carr A (2019), “Identification of human tendon cell populations in healthy and diseased tissue using combined single cell transcriptomics and proteomics,” BioRxIV, doi.org/10.1101/2019.12.09.869933.

Kim, D. H., Han, K., Gupta, K., Kwon, K. W., Suh, K. Y., and Levchenko, A. (2009), “Mechanosensitivity of fibroblast cell shape and movement to anisotropic substratum topography gradients,” Biomaterials, 30 (29), 5433–5444. DOI: 10.1016/j.biomaterials.2009.06.042.

Lee, C. H., Shin, H. J., Cho, I. H., Kang, Y. M., Kim, I. A., Park, K. D., and Shin, J. W. (2005), “Nanofiber alignment and direction of mechanical strain affect the ECM production of human ACL fibroblast,” Biomaterials, 26 (11), 1261–1270. DOI: 10.1016/j.biomaterials.2004.04.037.

Martins, J. A., Lach, A. A., Morris, H. L., Carr, A. J., and Mouthuy, P. A. (2020), “Polydioxanone implants: A systematic review on safety and performance in patients,” J Biomater Appl, 34 (7), 902–916. DOI: 10.1177/0885328219888841.

Mouthuy, P. A., Zargar, N., Hakimi, O., Lostis, E., and Carr, A. (2015), “Fabrication of continuous electrospun filaments with potential for use as medical fibres,” Biofabrication, 7 (2), 025006. DOI: 10.1088/1758-5090/7/2/025006.

Nezhentsev A, A. R., Baldwin M, Mimpen Y, Augustyniak E, Isaacs M, Mouthuy PA, Carr A, Snelling S (2021), “In vitro evaluation of the response of human tendon-derived stromal cells to a novel electrospun suture for tendon repair,” Translational Sports Medicine, 4, 409–418.

Riel, H., Lindstrom, C. F., Rathleff, M. S., Jensen, M. B., and Olesen, J. L. (2019), “Prevalence and incidence rate of lower-extremity tendinopathies in a Danish general practice: a registry-based study,” Bmc Musculoskeletal Disorders, 20. DOI: ARTN23910.1186/s12891-019-2629-6.

Smith, R. D., Carr, A., Dakin, S. G., Snelling, S. J., Yapp, C., and Hakimi, O. (2016), “The response of tenocytes to commercial scaffolds used for rotator cuff repair,” Eur Cell Mater, 31, 107–118. DOI: 10.22203/ecm.v031a08.

Still C, C. W., Sherman S, Sochacki K, Dragoo J, Qi L (2021), “Single-cell transcriptomic profiling reveals distinct mechanical responses between normal and diseased tendon progenitor cells,” Cell Reports Medicine, 2, 100343.

Stolk, M., Klatte-Schulz, F., Schmock, A., Minkwitz, S., Wildemann, B., and Seifert, M. (2017), “New insights into tenocyte-immune cell interplay in an in vitro model of inflammation,” Sci Rep, 7 (1), 9801. DOI: 10.1038/s41598-017-09875-x.

Tabula Muris, C., Overall, c., Logistical, c., Organ, c., processing, Library, p., sequencing, Computational data, a., Cell type, a., Writing, g., Supplemental text writing, g., and Principal, i. (2018), “Single-cell transcriptomics of 20 mouse organs creates a Tabula Muris,” Nature, 562 (7727), 367–372.

Yin, Z., Hu, J. J., Yang, L., Zheng, Z. F., An, C. R., Wu, B. B., Zhang, C., Shen, W. L., Liu, H. H., Chen, J. L., Heng, B. C., Guo, G. J., Chen, X., and Ouyang, H. W. (2016), “Single-cell analysis reveals a nestin(+) tendon stem/progenitor cell population with strong tenogenic potentiality,” Science Advances, 2 (11).

